# Single-cell transcriptomics elucidates *in vitro* reprogramming of human intestinal epithelium cultured in a physiodynamic gut-on-a-chip

**DOI:** 10.1101/2021.09.01.458444

**Authors:** Woojung Shin, Zhe Su, S. Stephen Yi, Hyun Jung Kim

**Affiliations:** Department of Biomedical Engineering, The University of Texas at Austin, Austin, TX 78712, USA; Department of Oncology, Livestrong Cancer Institutes, Dell Medical School, The University of Texas at Austin, Austin, TX 78712, USA; Oden Institute for Computational Engineering and Sciences (ICES), and Interdisciplinary Life Sciences Graduate Programs (ILSGP), The University of Texas at Austin, Austin, TX 78712, USA; Department of Medical Engineering, Yonsei University College of Medicine, 50-1 Yonsei-ro, Seodaemun-gu, Seoul, 03722 Republic of Korea; Wyss Institute for Biologically Inspired Engineering at Harvard University, Boston, MA 02115, USA

## Abstract

The microphysiological human gut-on-a-chip has demonstrated *in vivo*-relevant cellular fidelity of intestinal epithelium compared to its cultures in a static condition^1, 2^. Microfluidic control of morphogen gradients and mechanical cues robustly induced morphological histogenesis with villi-like three-dimensional (3D) microarchitecture, lineage-associated cytodifferentiation, and physiological functions of a human intestinal Caco-2 epithelium^3, 4^. However, transcriptomic dynamics that orchestrates morphological and functional reprogramming of the epithelium in a microphysiological culture remains elusive. Single-cell transcriptomic analysis revealed that a gut-on-a-chip culture that offers physiological motions and flow drives three distinctive subclusters that offer distinct gene expression and unique spatial representation in 3D epithelial layers. The pseudotemporal trajectory of individual cells visualized the evolutionary transition from ancestral genotypes in static cultures into more heterogeneous phenotypes in physiodynamic cultures on cell cycles, differentiation, and intestinal functions including digestion, absorption, drug transport, and metabolism of xenobiotics. Furthermore, the inversed transcriptomic signature of oncogenes and tumor-suppressor genes of Caco-2 cells verified that a gut-on-a-chip culture drives a postmitotic reprogramming of cancer-associated phenotypes. Thus, we discovered that a physiodynamic on-chip culture is necessary and sufficient for a cancer cell line to be reprogrammed to elicit *in vivo*-relevant heterogeneous cell populations with restored normal physiological signatures.

## Introduction

The Caco-2 human colon adenocarcinoma line has been extensively used in biomedical studies and considered as a gold standard for estimating intestinal permeability of drugs in pharmaceutical validation over the past decades^5-7^. The Caco-2 epithelium demonstrates enterocyte-like phenotype with transcriptomic signatures of normal small intestinal epithelium when cultured for 3 weeks in a static condition despite its cancer origin^8-10^. However, Caco-2 cells cultured as a 2D monolayer have shown a considerable deficiency in columnar 3D morphology, physiological differentiation, and basic intestinal functions including drug metabolism and mucus production^11^. Notably, epithelial cultures in a gut-on-a-chip significantly induced the cytodifferentiation and histogenesis of Caco-2 cells under physiodynamic fluid shear and mechanical deformations^1, 3, 4^. The biochemical gradient and mechanical cues reprogrammed the phenotypic outcomes including spontaneous 3D morphogenesis, lineage-dependent differentiation, mucus production, drug metabolism, and flow-dependent modulation of canonical Wnt signaling and subsequent subcellular responses. These observations strongly suggest that the phenotypic characteristics of Caco-2 cells are significantly perturbed by the microphysiological manipulation during cultures.

## Results and Discussion

To identify the underlying transcriptomic dynamics that lead to phenotypic reprogramming of the epithelium cultured in a gut-on-a-chip, we performed single-cell RNA sequencing (scRNA-seq) to compare the transcriptome profiles between on-chip and static Transwell cultures. Briefly, Caco-2 cells with the same passage number were grown in each culture format for approximately 7 days (Fig. 1a). In a Transwell, cells adherent on an extracellular matrix (ECM)-coated porous membrane formed a 2D monolayer with intact cell-cell junctions (Fig. 1a, Transwell). On the contrary, Caco-2 epithelium grown in a gut-on-a-chip experienced physiological flow (50 µL/h, equivalent shear stress at 0.02 dyne/cm^2^) and peristalsis-like rhythmical motions (10% in cell strain, 0.15 Hz in frequency), resulting in a recreation of villi-like 3D epithelial morphology as previously discovered (Fig. 1a, Gut-on-a-chip)^1, 3, 4^. Then, we performed scRNA-seq analyses (10x Genomics library preparation platform) followed by bioinformatic assessments including data quality control, variable gene selection, dimensionality reduction, and cell clustering using Seurat 3^12^. Differential gene expression from scRNA-seq results resolved four culture-dependent subclusters mapped on a Uniform Manifold Approximation and Projection (UMAP)^13^ (Fig. 1b). The physiologically dynamic culture in a gut-on-a-chip was the only source of variations into three clusters (CHIP1 - 3), whereas the static Transwell culture resulted in an isolated cluster (TW1). Interestingly, a heatmap that plots the top 15 genes that were most significantly and differentially expressed in each cluster involved the genes that regulate intestinal epithelial functions such as proliferation, regeneration, cell cycle, and other differentiated functions including molecular transport and drug metabolism (Fig. 1c). For instance, CHIP1 cluster highlights genes pertinent to the regulation of cell cycle and growth (*CDK1, AREG*, and *UBE2C*) and mucosal barrier function (*MUC13*). CHIP2 shows significant upregulation of genes associated with cell division and mitosis (*PTTG1, CDC20, CCNB1, DDX21*, and *MYC*). In CHIP3, genes related to molecular transport (*SLC2A3, SLC6A8, FXYD3, SLC11A2*, and *APOA4*) are highly and distinctively expressed. TW1 cluster, in contrast, includes upregulated genes related to cell mitosis (*HEPACAM2*), mitogens (*FGB, FGG, FGA*), and a colorectal cancer marker (*PHGR1*)^14^. It is noted that the mitotic genes (*CDK1* and *UBE2C*) upregulated in CHIP1 were significantly downregulated in CHIP2, suggesting that heterogeneous cellular reprogramming occurred in individual clusters of cells in the gut-on-a-chip culture.

**Fig. 1.**
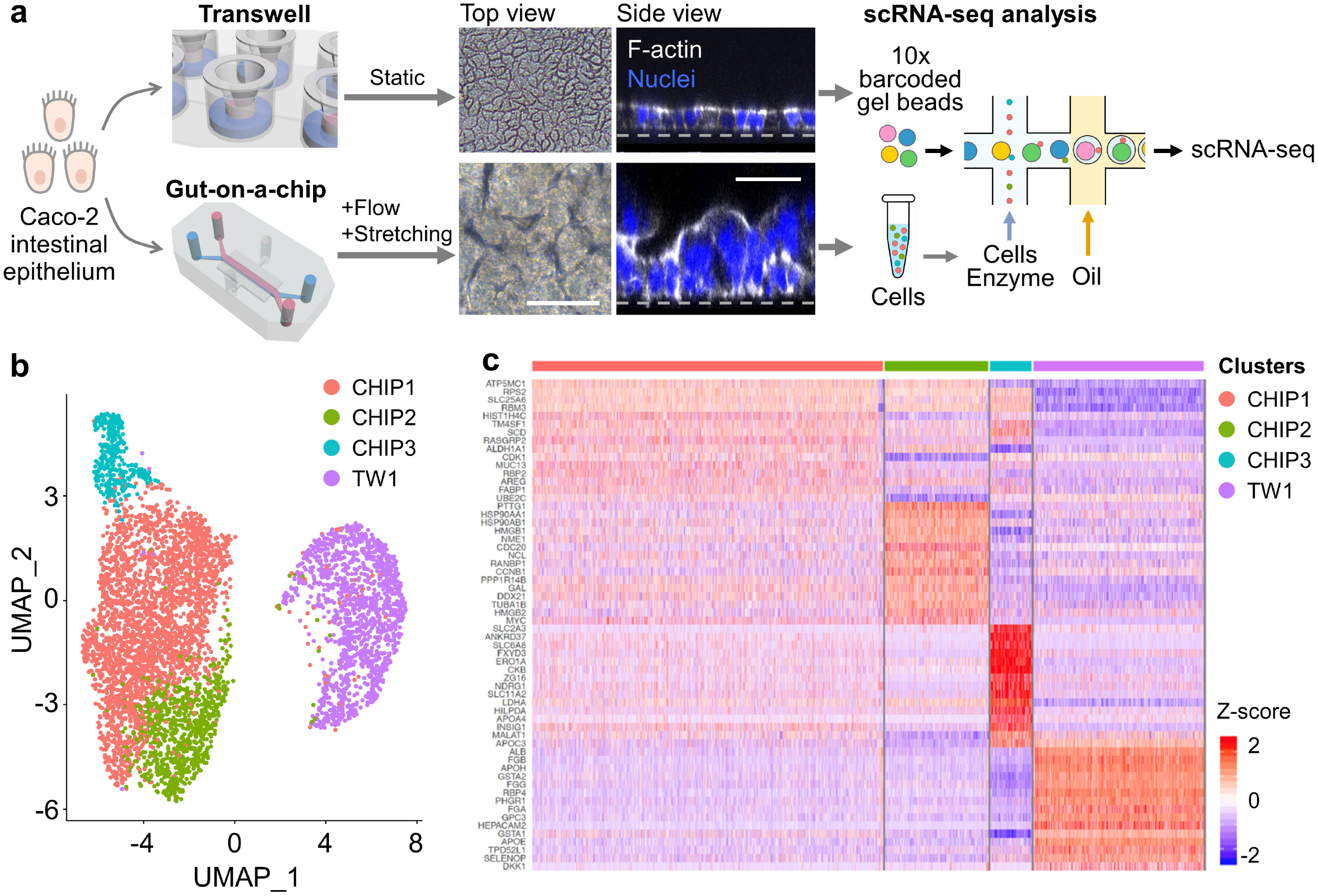
Single-cell characterization of culture-dependent epithelial heterogeneity. **a**, An overview of scRNA-seq analysis of Caco-2 cells cultured in either a static Transwell or a physiodynamic gut-on-a-chip. Caco-2 cells cultured in a Transwell insert formed a 2D monolayer, whereas the same cells cultured in a gut-on-a-chip under flow and stretching motions regenerated a 3D villi-like epithelial microstructure (Top view, phase contrast micrographs; Side view, immunofluorescence micrographs; dashed lines, location of the basement porous membrane). Cultured cells were harvested and processed for scRNA-seq analysis by the 10x Genomics platform. **b**, A non-linear dimensionality reduction displayed in a UMAP that plots single cells from both culture formats. Cells from the Transwell culture have a homogeneous population (TW1), whereas the gut-on-a-chip culture demonstrated three heterogenous subclusters (CHIP1 - 3). **c**, A heatmap showing the top 15 genes differentially expressed in each subcluster. A complete list of selected genes is provided in Extended Data Table 1. Bars, 50 µm.

Based on primary clustering, we retroactively tested if each cell cluster may encode spatial information in the 3D epithelial layer because the gut-on-a-chip culture allowed 3D regenerative histogenesis that contains individual cell subclusters. We visually verified this hypothesis in the established histogenic structures in a gut-on-a-chip. Among the differentially expressed genes, we chose *MUC13* and *CDK1* (CHIP1), *CCNB1* (CHIP2), and *SLC11A2* and *SLC6A8* (CHIP3) that the expression in one cluster is significantly higher than the others (Fig. 2a and 2b) and performed immunofluorescence imaging by targeting encoded proteins. The 3D rendered views revealed that the fluorescence signals of each marker uniquely illustrated the localized niche in the 3D epithelial layers recreated in the gut-on-a-chip (Fig. 2c and 2d). We found that the spatial signal of MUC13 and CDK1 (CHIP1) and CCNB1 (CHIP2) was predominantly localized in the middle-lower region, whereas a considerable number of cells expressing SLC11A2 and SLC6A8 (CHIP3) are positioned in the upper-middle region of the 3D epithelial layer. This observation suggested that the clusters derived from a gut-on-a-chip culture may undergo distinct spatial organization of proliferative and differentiating cells along the migration axis from the basement membrane toward the upper region of the mucosal surface, reminiscent of the crypt-villus axis *in vivo*. We previously reported a similar epithelial behavior using a gut-on-a-chip, where EdU-positive proliferative cells migrated from the basal region to the epithelial tip^3^. The result was also, in part, correlated to the physiological functions of genes that we highlighted in this analysis. For instance, *CDK1*, representing CHIP1, is involved in the cell cycle and apoptotic regulations^15^, and the higher expression in the proliferating basal area is concordant with this function. CHIP2 representing *CCNB1* that encodes cyclin B1 regulates the transition process from G2 to M phase in cell cycle^16^. Therefore, the localized expression of *CCNB1* in the middle-lower region of the 3D epithelial layer agrees with its known roles and localization. The selected genes for CHIP3, *SLC6A8* and *SLC11A2* are a Na^+^ Cl^-^ dependent creatine transporter and a metal transporter^17, 18^, respectively. These transporters are known to be expressed predominantly in the intestinal villi rather than the basal crypt *in vivo*, which was visually confirmed.

**Fig. 2.**
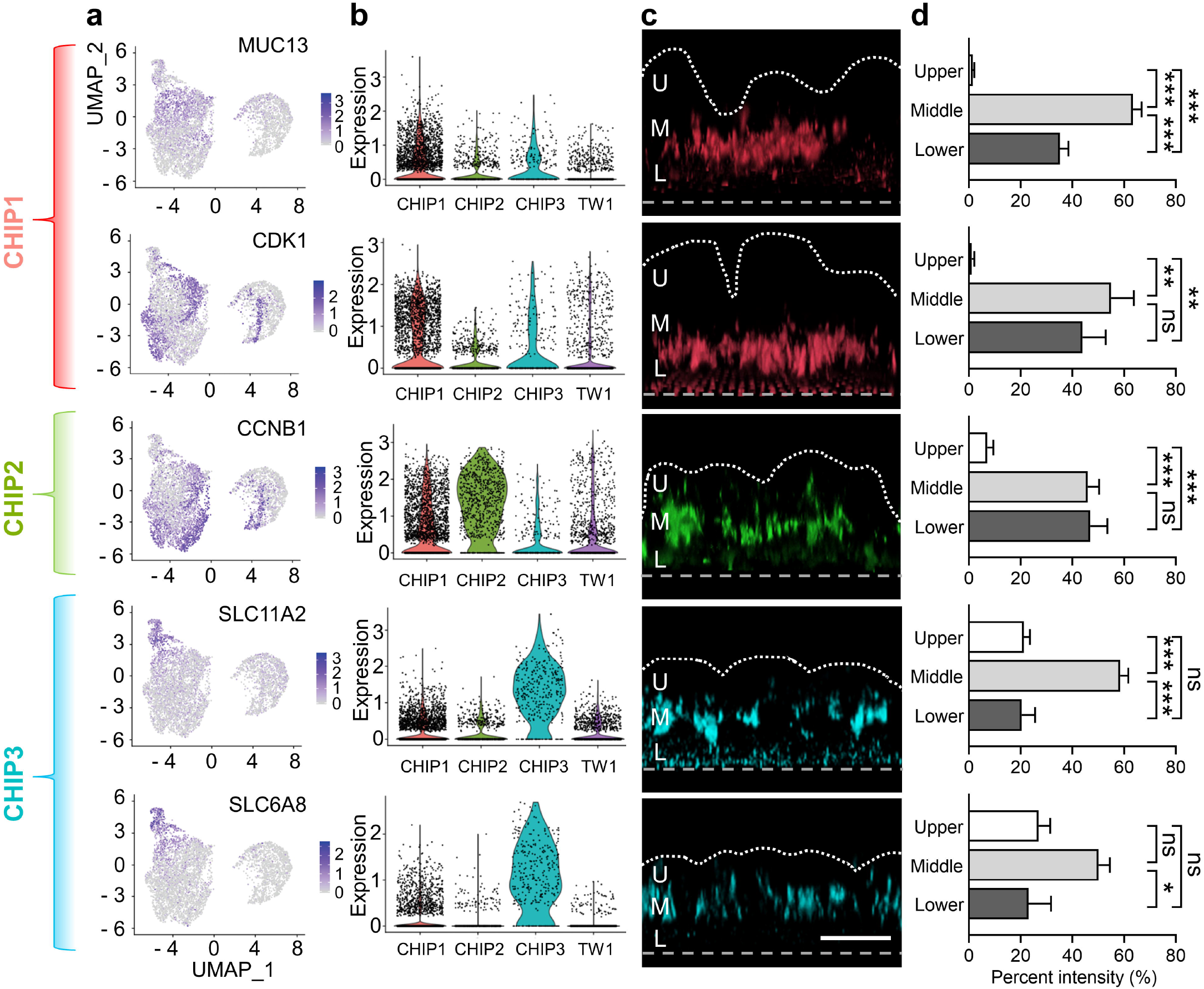
Spatial mapping of translated gene products after physiodynamic cellular reprogramming. **a**, Expression of 5 selected gene products that represent each CHIP subcluster is highlighted in the UMAP. **b**, Violin plots corresponding to the selected genes in **a. c**, The spatial localization of the selected genes in 3D reconstruction views visualized by immunofluorescence confocal microscopy. Bar, 100 µm. White dotted lines indicate the outline of the 3D epithelial layers. Dashed lines represent the location of porous membranes. U, upper, M, middle, and L, lower region. **d**, Quantifications of fluorescent signals in spatially resolved regions along the vertical direction. N=4, ns, not significant. \**P*<0.05, \*\**P*<0.001, \*\*\**P*<0.0001. Error bars, standard error of mean.

The selected top 15 genes (Fig. 1c) and the spatial expression of representative genes in each cluster (Fig. 2) suggested that CHIP3 contains the most differentiated cell population, whereas CHIP1 and CHIP2 are dominated by proliferative cells that are transient to a differentiated stage. This cellular and morphological reprogramming process was additionally characterized by an unbiased pseudotime trajectory analysis to dissect the evolutionary transition of the cells and their clusters depending on the culture formats (Fig. 3a). The merged 2D trajectory of all the clusters outlined a pseudotemporal propagation from the ancestral heredity (i.e., pseudotime is zero; dark blue) toward bifurcated branches (i.e., bright blue). When we set TW1 as the most ancestral position (e.g., the left end branch), the cells in CHIP1 showed the largest spectrum of pseudotemporal distribution. On the contrary, the pseudotemporal points of cells in CHIP2 and CHIP3 are found at the farthest positions from the ancestral TW1, suggesting that the cells cultured in the gut-on-a-chip were significantly evolved in pseudotime configuration than in the static Transwell culture. This result is strongly correlated with the identified spatial information of cell clusters verified in Fig. 2.

**Fig. 3.**
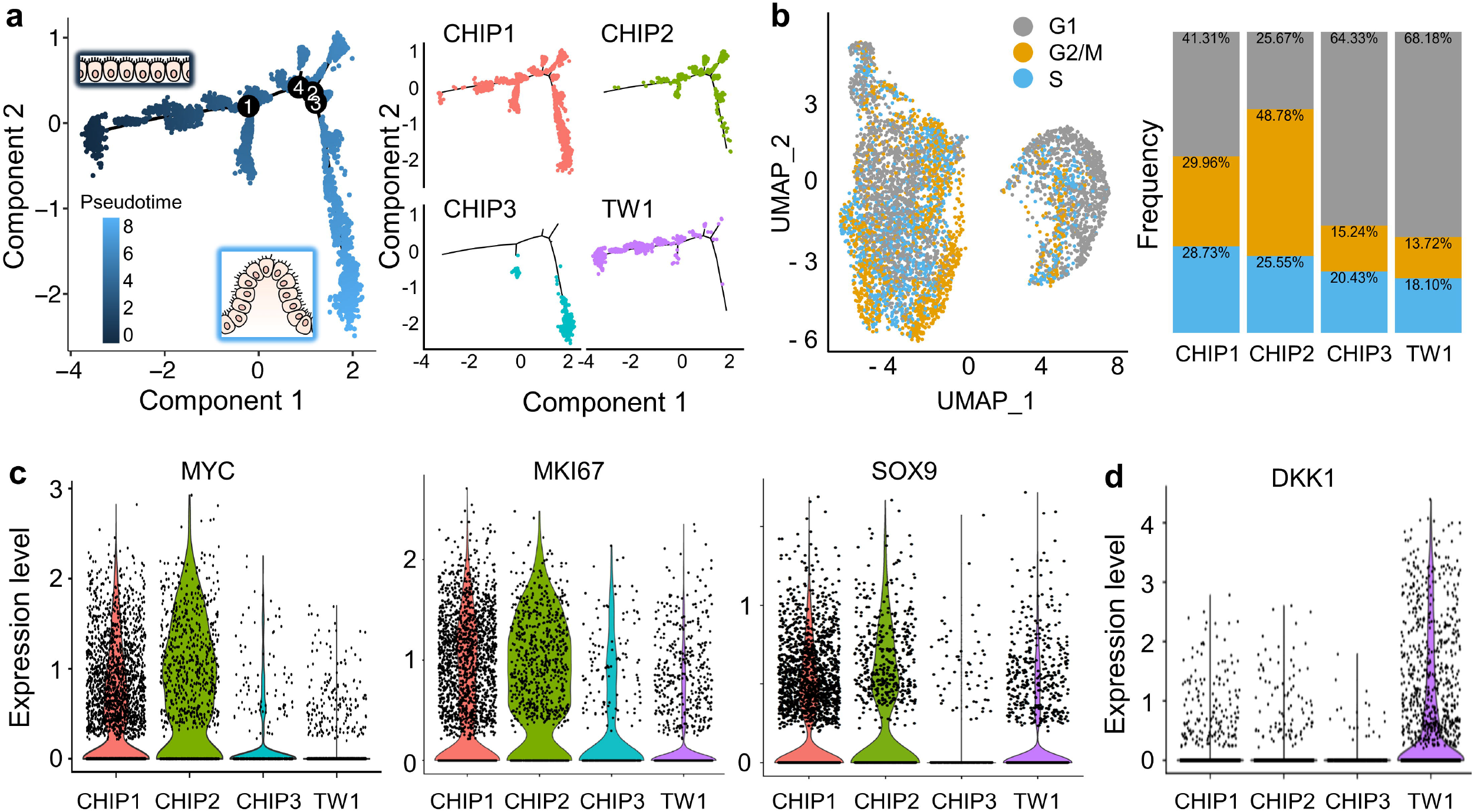
Heterogenous reprogramming of cell cycle and developmental process. **a**, A collective pseudotemporal trajectory of single cells cultured in either a Transwell or a gut-on-a-chip (left) and the individual pseudotime with a cluster-dependent resolution (right). (Left) The cells in TW1 cluster (dark blue) were assigned as the ancestral pseudotime, in which the cells in CHIP clusters were widely distributed in branched pseudotemporal trajectories (lighter blue), indicating the heterogeneous evolutionary reprogramming. The branching points are labeled in circled numbers. The inset schematics illustrate the shape of a cell layer in TW1 (left top; 2D) or CHIP (right bottom; 3D). (Right) The pseudotemporal distribution of individual subclusters. **b**, Distribution of individual single cells that show distinct cell cycles in G1 (grey), G2/M (orange), or S phase (sky blue) plotted on the UMAP (left) and the frequency of single cells in each subcluster. **c & d**, Violin plots that represent the significant upregulation of **c**, *MYC, MKI67*, and *SOX9* in CHIP1 and CHIP2 clusters and **d**, an elevated expression of *DKK1* in TW1 cluster.

The cell cycle analysis displayed on the UMAP (Fig. 3b, left) reveals that the subclustered cells in CHIP1 and CHIP2 cultured in a gut-on-a-chip showed substantially higher populations of G2/M- and S-phases than the cells grown in a Transwell (Fig. 3b, right). The cells in TW1 cluster were composed of the largest G1 phase (68.18%) and the least G2/M (13.72%) and S phase (18.10%). Interestingly, the cell cycle frequency in CHIP3 cluster (64.33, 15.24, and 20.43% of G1, G2M, and S phases, respectively) showed the highest similarity to the frequency in TW1, suggesting that the cellular dynamics of proliferation characterized by the cell cycle analysis is minimal in both clusters. Regardless of the similar cell cycle patterns, it is noted that the morphological and spatial transcriptomic signatures of the Caco-2 cells cultured in a gut-on-a-chip were considerably distinct from the ones prepared in a Transwell, indicating that the microphysiological cultures dramatically induce cellular reprogramming of an immortalized cell line in shapes and functions with greater *in vivo* relevance. The profile of mitochondrial genes (%MT) also showed a good agreement with the cell cycle analysis (Fig. S1)^19, 20^. Indeed, both CHIP1 and CHIP2 clusters showed a significantly elevated population of cells (<75%) in G2/M or S phase than CHIP3, suggesting that the dynamics of the cell cycle and mitosis can be considerably heterogeneous when cells are cultured in a microphysiological culture device.

It is notable that the heterogeneity of transcriptomic signature regarding cell proliferation was also represented in the expression of genes encoding proto-oncogenic transcription factor (*MYC*), proliferation (*MKI67*), and stem cell (*SOX9*), in which both CHIP1 and CHIP2 showed the remarkable elevation of those genes than CHIP3 and TW1 clusters (Fig. 3c). Interestingly, although CHIP3 cluster showed similar patterns in cell cycle and proliferation with TW1, a representative Wnt-antagonizing gene, Dickkopf-related protein 1 (*DKK1*), that is critical for intestinal morphogenesis in a gut-on-a-chip^4^ was minimally expressed in all the CHIP clusters, whereas TW1 was a dominant cluster that shows significant upregulation of *DKK1* (Fig. 3d). This result shows good accordance with our previous finding that the removal of basolaterally secreted DKK1 molecules leads to the 3D morphogenesis of intestinal epithelial cells, whereas a static Transwell culture that accumulates extracellular DKK-1 considerably restricts the morphogenic process regardless of the culture period^3^. The transcriptomic heterogeneity of epithelial growth dynamics was also verified by a couple of gene ontology (GO) snapshots on the heterogeneous signaling perturbations of Wnt (Fig. S2), Notch (Fig. S3), and bone morphogenetic protein (BMP) pathways (Fig. S4) as well as planar cell polarity (Fig. S5).

Next, we examined the heterogeneity of transcriptomes in each cluster related to fundamental epithelial functions such as nutrient absorption and transport, biosynthetic process, and drug metabolism. Interestingly, we found that the expression of genes germane to monosaccharide transport, sugar transmembrane transporter activity, carbohydrate transporter activity, and sodium channel regulator activity (Fig. 4a), glucose catabolic process, and fatty acid biosynthesis (Fig. 4b) were all significantly higher in CHIP3 than the other clusters. This result is well matched with the spatial information that the cells in CHIP3 are located in the upper layer of the intestinal epithelium because these transport activities, catabolism, and fatty acid biosynthesis are regulated at the mucosal surface by enterocytes^21^. The gene set enrichment analysis (GSEA) that compares the most significant 33 GO terms also verified that the gene sets associated with intestinal molecular transport were strongly upregulated in CHIP3 (Fig. S6). We additionally mapped the overall transcriptome profile of transporters targeting water, vitamin, bile salt, lipid, organic solute, inorganic solute, amino acid, nucleotide, metal ion, and sugar, confirming heterogeneous reprogrammed features of Caco-2 cells cultured in a gut-on-a-chip (Fig. S7). Importantly, approximately two-thirds of the genes linked to GO Drug Transport process were considerably upregulated in CHIP clusters (e.g., *SLC1A5, RALBP1, GAL, TCN2, SYT8*, and *SLC25A5*) while the genes highlighted in TW1 cluster represented conserved epithelial functions such as glutamate transport (*SLC1A1*) or proton-coupled peptide absorption (*CA2*). Moreover, the gut-on-a-chip culture elicited Caco-2 cells to show reprogrammed drug metabolizing characteristics, where the heterogeneous population of single cells harvested from the gut-on-a-chip showed elevated expression of genes encoding Phase I (*CYP1A1, CYP2S1*, and *ALDH1A1*; Figs. 4d and S8) and Phase II (*AHCY*) enzymes as well as a Phase III transporter (*SLC6A8*; Fig. 4e)^22^.

**Fig. 4.**
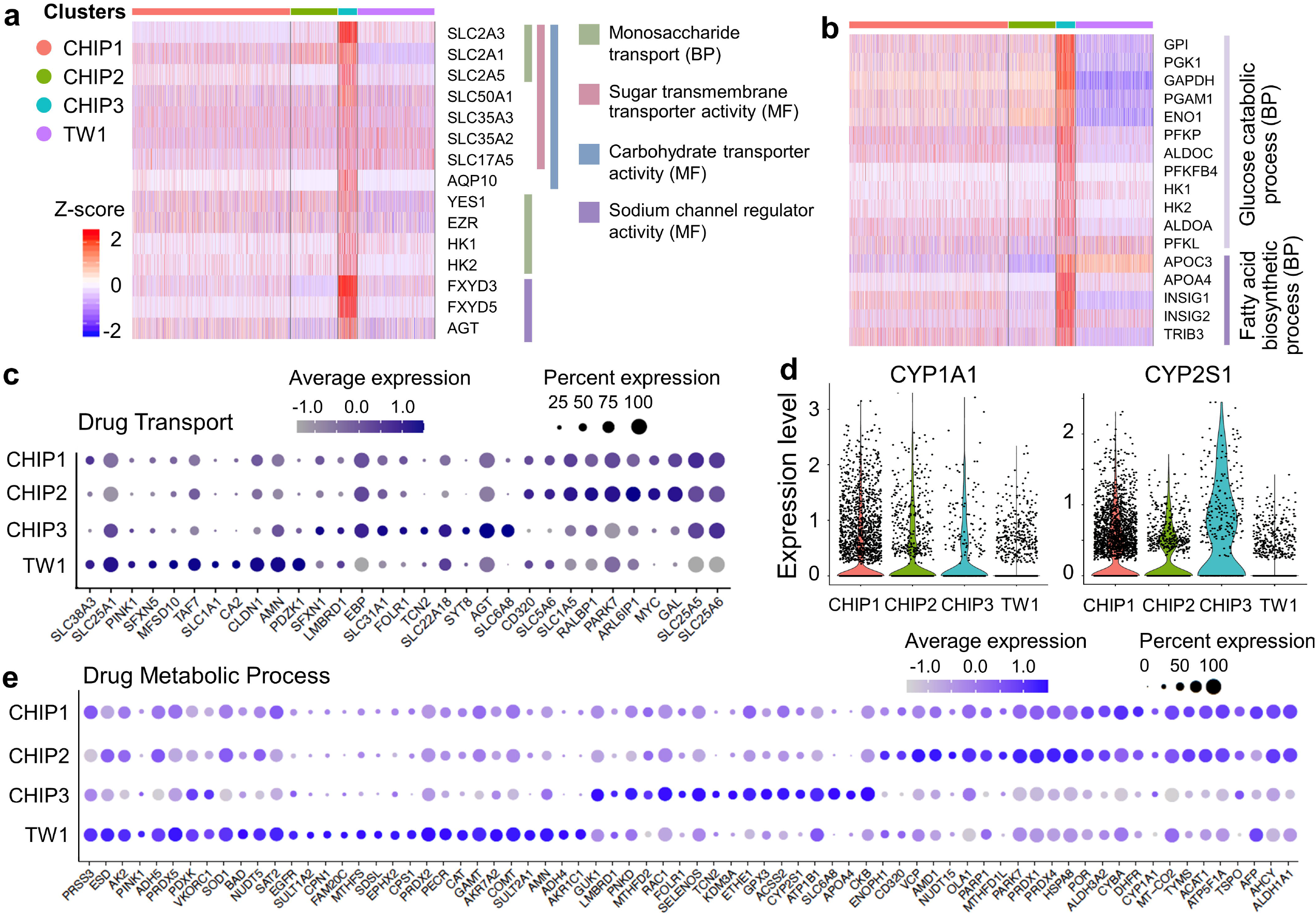
Reprogramming on intestinal epithelial functions and drug metabolism. **a**, A heatmap that summarizes the expression of genes associated with the GO terms of monosaccharide transport (BP), sugar transmembrane transporter activity (MF), carbohydrate transporter activity (MF), and sodium channel regulator activity (MF). **b**, A heatmap that maps the expression of genes involved in the GO terms of glucose catabolic process (BP) and fatty acid biosynthetic process (BP). BP, biological process, MF, molecular function. **c**, A dot plot that displays the reprogrammed single-cell transcriptomes pertinent to drug transport. **d**, Violin plots that highlight the elevated expression of *CYP1A1* and *CYP2S1* in CHIP clusters compared to TW1. **e**, A dot plot that visualizes the reprogrammed expression of genes regulating drug metabolic process.

Surprisingly, Caco-2 cells that underwent the on-chip microphysiological growth showed a significant transcriptomic perturbation of genes germane to oncogenic or tumor-suppressive functions. Using the Cancer Gene Census database^23^, we selected out genes associated to colorectal cancer and grouped them into oncogenes and tumor suppressors. The oncogenes revealed the diminished expression in all the CHIP clusters compared to TW1 cluster (Fig. 5a). On the contrary, the tumor-suppressor genes showed substantial increases in all of the CHIP clusters, especially in CHIP3, compared to TW1 (Fig. 5b). We confirmed that some colon-specific cancer stem cell markers including *PHGR1, PROM1* (encoding CD133), and *DPP4* also showed significant reduction of expressions (Fig. 5c), suggesting that a physiodynamic gut-on-a-chip culture of a cancer cell line drives a postmitotic reprogramming of cancer-associated phenotypes along with other essential epithelial functions.

**Fig. 5.**
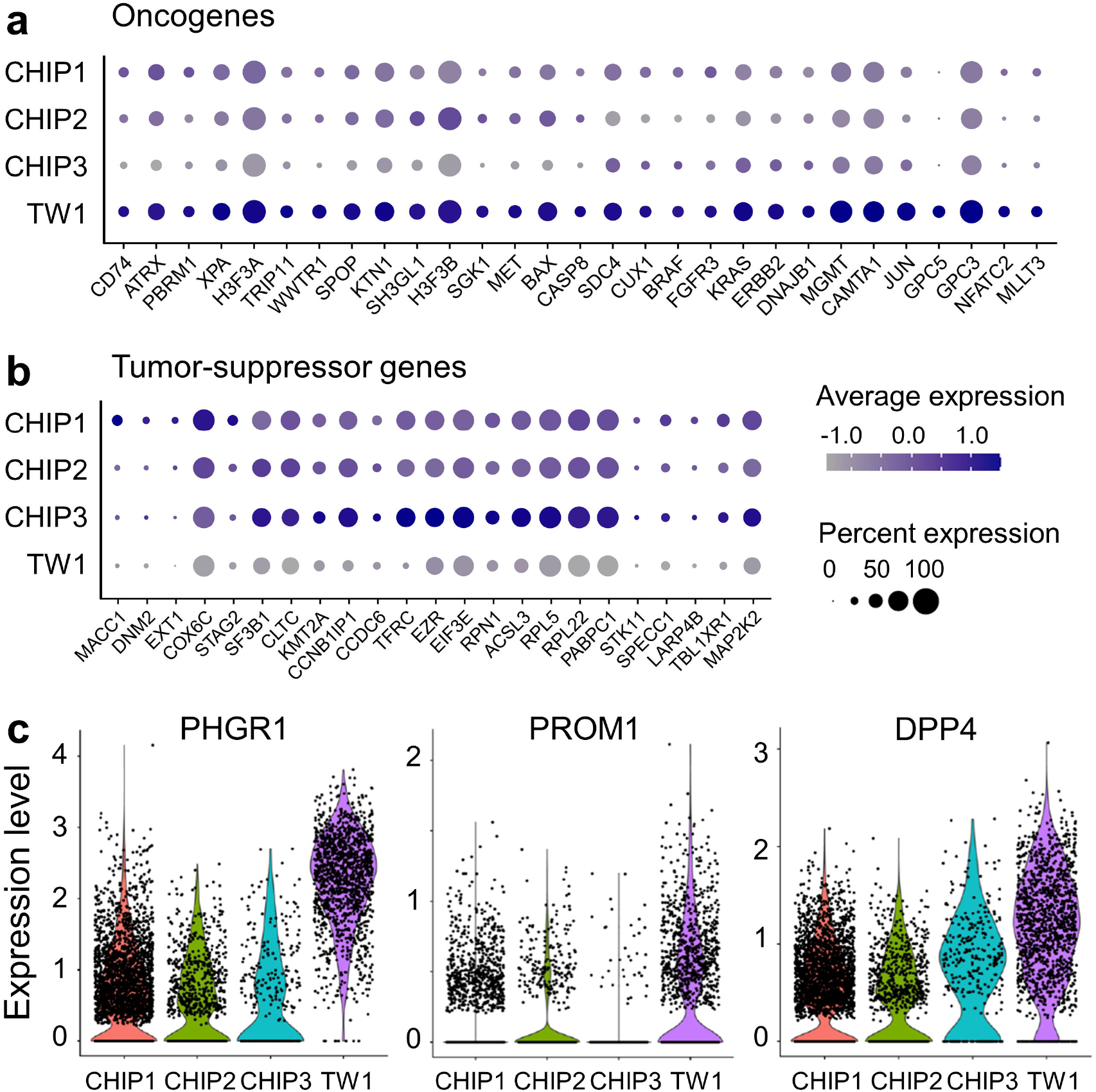
Reprogrammed transcriptomic feature of cancer inheritance on Caco-2 cells. Dot plots that summarize the reprogrammed transcriptomic signature of **a**, oncogenes and **b**, tumor-suppressor genes. **c**, Violin plots that visualize the expression of representative markers of colon cancer stem cells, *PHGR1, PROM1*, and *DPP4* between subclusters.

Our study presents a number of impacts. First, this study did not involve any cellular and molecular engineering (e.g., CRISPR-based genome engineering) nor surrounding cellular components (e.g., stromal cells, immune cells, or microbiome) that can potentially perturb cellular plasticity at various disease milieus. Hereby, we confirmed that a physiodynamic gut-on-a-chip culture is necessary and sufficient for a cancer cell line to be reprogrammed to elicit *in vivo*-relevant heterogeneous physiological functions. Second, the reversion of cancerous characteristics to the normal by simply culturing in a gut-on-a-chip suggests a novel avenue of mechanistic cancer studies. Our results reveal that conventional static cultures, from immortalized cell lines to patient-derived organoid cultures, may need to be proactively evolved toward physiodynamic cultures because *in vitro* static cultures neither restore the *in vivo*-relevant original phenotypic characteristics nor recapitulate physiological and pathological responses. Indeed, the microphysiological human gut-on-a-chip may repurpose a new platform to accurately manipulate the tissue microenvironment to investigate cellular developmental process, molecular interactions in a spatiotemporal landscape, selected stimulation with a directional configuration (e.g., apical vs. basolateral), and time-resolved multi-modal assessments. Finally, the legitimacy and validity of Caco-2 cells in pharmaceutical applications will be considerably repurposed when the cells are reprogrammed and functionalized by a gut-on-a-chip culture. Since Caco-2 cells have been used in pharmaceutical and biomedical studies over the past couple of decades, it may be a commercially attractive option to induce directional reprogramming of Caco-2 cells and enhance the utility for defined applications.

In summary, we elucidated that the physiodynamic culture in a human gut-on-a-chip enhances intestinal physiological functions of Caco-2 cells, where notably, chemical and the nutritional environment was identical to the static Transwell cultures. We verified that Caco-2 cells spontaneously undergo reprogramming of transcriptomic signature, pseudotemporal evolution, spatial reorganization of epithelium in a basal-mucosal axis, restoration of original intestinal functions, and reversion from a cancerous to a normal phenotype when cultured in a gut-on-a-chip. We foresee that our proof-of-principle study will disseminate a practical and implementable protocol to be applicable to other tissue-derived or iPSC-derived cells under a defined biomechanical, chemical, and physiological condition.

