## Supplementary information for "Single-cell transcriptomics elucidates *in vitro* reprogramming of human intestinal epithelium cultured in a physiodynamic gut-on-a-chip"

Or

S. Stephen Yi, PhD
Department of Oncology, Livestrong Cancer Institutes, Dell Medical School
The University of Texas at Austin


### Methods

#### Microfabrication of a gut-on-a-chip

A gut-on-a-chip device was fabricated using polydimethylsiloxane (PDMS; SYLGARD 184 silicone elastomer kit, Dow Corning) by following the soft lithography method that was previously established[1, 2]. Briefly, silicon molds that are patterned with SU-8 (Microchem) with the designs of upper and lower microchannels were prepared via a photolithography method[3, 4]. A silicon mold that has an array of pillars (10 μm in diameter, 25 μm in height, and 25 μm in spacing) was used to fabricate a sheet of a porous membrane. Once the clean silicon molds for upper, lower, and membrane layers are prepared, precured PDMS solution was prepared by vigorously mixing base silicone polymer and curing agent (mass ratio, 15:1) and degassing it in a vacuum desiccator at room temperature (RT) for 20 min. Degassed PDMS mix was poured on the silicon molds with target thicknesses of 7 mm and 1 mm for making an upper and a lower PDMS layer, respectively, then cured in a dry oven at 60°C for 12 h. To fabricate a porous membrane layer, ~1 mL of uncured PDMS solution was poured per mold, spread on the mold, and covered with a sheet of fluoropolymer-coated polyester film (3M Scotchpak Release Liner, 3M), then pressed with a weight (3 kg) while curing in a dry oven at 60°C for 12 h. After punching holes (diameter, 2 mm) in an upper layer for connecting silicone tubing, the upper layer was sequentially bonded to a membrane layer followed by a lower microchannel layer using a handheld corona treater (Electro-Technic). A complete gut-on-a-chip device was connected with silicon tubing (Tygon 3350), sterilized with 70% ethanol, and dried in a dry oven at 60°C for overnight until use.

#### Culture of Caco-2 cells in a Transwell and a gut-on-a-chip

Caco-2 BBE line (Harvard Digestive Disease Center) used in this study was routinely cultured in T75 culture flasks using Dulbecco's Modified Eagle medium (DMEM; Gibco) supplemented with 20% (v/v) fetal bovine serum (FBS, heat inactivated; Gibco) and antibiotics (100 U/mL penicillin, 100 μg/mL streptomycin, final concentration; Gibco). To culture cells in a Transwell insert, dissociated Caco-2 cells (final density, $1\times10^6$ cells/mL) were seeded in the porous insert (0.4 μm in pores; Corning) pre-coated with a mixture of extracellular matrix (ECM; 100× diluted Matrigel (Corning) and 30 μg/mL of collagen type I (Gibco)) and incubated in a humidified $CO_2$ incubator at 37°C by changing culture medium every other day up to 5 days. To culture Caco-2 cells in a gut-on-a-chip, the surface of microchannels in a fabricated device was activated with ultraviolet/ozone treatment for 40 min (UVO, Zelight Company Inc.), then introduced with the ECM mixture (Matrigel and collagen I) in a humidified $CO_2$ incubator at 37°C for 1 h. After the ECM solution in the microchannels was replaced with a serum-containing culture medium, disassociated Caco-2 cells (final density, $1\times10^7$ cells/mL) were seeded in the upper microchannel, and incubated in a humidified $CO_2$ incubator at 37°C for 1 h for cell attachment. Unbound cells were washed by flowing the culture medium and flow in the upper channel was maintained (volumetric flow rate at 30 μL/h) until the cells form a confluent monolayer (24 – 36 h). The culture was further maintained under flow in both upper and lower microchannels at a flow rate of 50 μL/h and rhythmical stretching motions (10% strain, 0.15 Hz frequency) for 5 days.

**Cell preparation for scRNA-seq analysis**

To harvest cells cultured in the Transwell, three inserts were washed with phosphate buffered saline (PBS, $Ca^{2+}$- and $Mg^{2+}$-free; Gibco) and treated with 200 μL of trypsin/EDTA (Gibco) for cell detachment at 37°C for 5 min. Cells were collected in a 15 mL conical tube and 5 mL of the cell culture medium was added to inactivate trypsin. The cell suspension was filtered through a 20 μm cell strainer (pluriSelect) to remove cell clumps, and centrifuged at 300×*g* at 4°C for 3 min. The cell pellet was then frozen in 500 μL of a freezing medium (10%, v/v, dimethyl sulfoxide in FBS). To harvest cells from the gut-on-a-chip culture for single-cell RNA-seq, the channels were washed with 500 μL of PBS, and incubated with 300 μL of trypsin/EDTA at 37°C for 5 min. Dissociated cell suspension was collected in a 15 mL conical tube and incubated with 5 mL of the cell culture medium for trypsin inactivation. Cells were centrifuged at 300×*g* at 4°C for 3 min and the cells were stocked in 500 μL of the freezing medium after aspiration of the supernatant. For library preparation, the frozen cells were thawed and resuspended in 10 mL of 2% (w/v) bovine serum albumin (BSA; MP Biomedicals) dissolved in PBS. Cells were then washed by repeating centrifugation (300×*g,* 4°C, for 3 min), aspiration, and resuspension (in 500 μL of 2% BSA solution). Cell suspensions were run through the Chromium (10x Genomic) using the Single Cell 3′ Reagent Kits v3, following the manufacturer's protocol. The library prepared samples were run for sequencing on the Illumina HiSeq Series with 50,000 target reads per cell.

**Image analysis**

Cell morphologies were monitored by phase contrast microscopy (DMi1, Leica Microsystems). To visualize the contour of cell microarchitecture formed in a Transwell insert and a gut-on-a-chip (Fig. 1a), we fixed the cells with 4% (w/v) paraformaldehyde (PFA; Alfa Aesar) for 10 min, permeabilized with 0.3% (v/v) Triton X-100 (VWR International) for 10 min, and stained with a mixture of CruzFluor 647 conjugated phalloidin (for F-actin; Santa Cruz Biotechnology) and

4’,6-diamidino-2-phenylindole dihydrochloride (DAPI for nuclei; final concentration, 1 μg/mL; Thermo Scientific) for 30 min, sequentially at room temperature under light protection. After each step, cells were rinsed with PBS to remove residual reagents.

To perform antibody-based immunofluorescence staining, cells were fixed (4% PFA, 10 min), permeabilized (0.3% Triton X-100 in 2% (w/v) BSA, 30 min), and blocked (2% BSA, 1 h) at room temperature. Between steps, cells were washed with PBS. Primary antibody solutions (mouse monoclonal anti-MUC13, 10 μg/mL, Abcam; rabbit monoclonal anti-CDK1, 1:50 dilution, Abcam; rabbit polyclonal anti-Cyclin B1, 1:100 dilution, Cell Signaling Technology; rabbit polyclonal anti-DMT1, 4 μg/mL, Abcam; rabbit polyclonal SLC6A8, 1:50 dilution, ThermoFisher Scientific) were incubated with the cells at room temperature for 3 h. After washing the cells with PBS, secondary antibody solutions (DyLight 488-conjugated goat anti-mouse antibody, Abcam; Alexa Fluor 555-conjugated goat anti-mouse antibody, Abcam) were incubated with the cells at room temperature in dark for 3 h. A mixture of CruzFluor 647 conjugated phalloidin (1× concentration) and DAPI (1 μg/mL, final concentration) was subsequently introduced for the counter staining of F-actin and nuclei, respectively, at room temperature for 30 min. Fluorescence imaging analysis was performed via a laser-scanning confocal microscope (DMi8, Leica Microsystems) using a 25× objective (NA 0.95, water immersion, Leica) coupled with the TCS SPE confocal system with solid state excitation laser sources of 405 nm, 488 nm, 532 nm, and 635 nm and an Ultra high dynamic PMT detector. To investigate the spatial expression in a gut-on-a-chip, z-stacked images were 3D reconstructed using a 3D module on LAS X (Leica Microsystems).

To analyze spatially localized fluorescence signal of the epithelial layers formed in gut-on-a-chip devices (Fig. 2d), the height of each epithelium was measured from the z-stacked images. The measured heights were divided into three sections (upper, middle, and lower) and the intensity of each section was quantified using ImageJ. The measured intensity was transformed into the percent intensity.

**Mapping and aggregation**

The sequence read files were imported to Cell Ranger count (v3.0, 10x Genomics) to align to the human reference and generate the unique molecular identifier (UMI) counts of gene-cell matrix by sample. After a quality control, 3,920 and 1,348 cells were captured from the gut-on-a-chip and the Transwell, respectively. Then Cell Ranger 'aggr' were used to aggregate the UMI counts matrixes of the two samples with batch effect correction[5].

**Quality control and transcriptomic analyses**

Using Seurat 3.0, firstly, the quality control was performed on Cell Ranger filtered counts. The cells and genes meeting the following conditions were kept: 1) the gene was detected at least in 3 cells; 2) the cell contained more than 2,000 and less than 6,000 detectable genes; 3) for a cell, the percentage of total mitochondrial gene UMI was less than 20%. Then, a global-scaling normalization ('logNormalize', scale factor is 10,000 by default) was performed and 200 highly variable features were listed. After the cell cycle scoring2 and regression, and scaling the data, the variable features were used for PCA analysis[6].

The number of principal components was determined using a gene correlation matrix by the 'elbow' method. Based on a Shared Nearest Neighbor (SNN) method, the cells were classified into distinct clusters on the dimensional reduction plots. Finally, the UMAP[7] was used for non-linear dimensional reduction, to reveal transcriptional characteristics of distinct cellular clusters. For cell clustering and non-linear dimensional reduction, 13 statistically significant PC dimensions were used.

**Trajectory analysis**

The normalized and log transformed gene-cell matrix from Seurat processed data was imported to Monocle 2. The top 10 differential expressed genes of all the 4 cell clusters calculated by 'FindAllMakers' in Seurat were collected, and then the genes which contained the GO term 'GO_CELL_CYCLE' were filtered out, then the rest were used as the ordering genes for cell trajectory analysis. Cells of TW1 were set as the root – time zero[8].

**Identification of marker genes and gene set enrichment analysis**

'FindMarkers' in Seurat 3.0 was used for analyzing the average log fold changes between two cell clusters. Ranking the log fold changes of all expressed genes, the gene set enrichment analysis (GSEA) was performed with C5 ontology gene sets (GO items) for any two groups - CHIP3 versus CHIP1,2[9].

**Data availability**

Data generated for this study will be available through the Gene Expression Omnibus (GEO). Online Content Methods, along with any additional Extended Data display items, are available in the online version of the paper.

**Statistical analysis**

Statistical analysis was performed in GraphPad Prism 9. To evaluate the statistical significance between groups, ordinary one-way ANOVA tests with multiple comparisons was performed (Fig. 2d).

**Acknowledgments**

This work was supported in part by the National Cancer Institute of the National Institutes of Health (NIH/NCI) F99/K00 Predoctoral to Postdoctoral Transition Award (K00CA245801 to W.S.), Asan Foundation Biomedical Science Scholarship (W.S.), Mogam Science Scholarship (W.S.), the National Institute of General Medical Sciences (NIH/NIGMS) Maximizing Investigators' Research Award (MIRA) (R35 GM133658 to S.S.Y.), Technology Impact Award of the Cancer Research Institute (UTA18-000889 to H.J.K.), and the Leona M. & Harry B. Helmsley Charitable Trust (Grant #1912-03604 to H.J.K.).

**Author Contributions**

W.S. and H.J.K. conceived and designed the study and performed experiments. S.S.Y. and H.J.K. supervised the overall research and provided intellectual input. W.S., Z.S., S.S.Y., and H.J.K. analyzed the data and wrote the manuscript.

**Competing interests**

The authors declare no conflict of interest.

**Extended Data**

**Extended Data Table 1. A list of the top 15 genes differentially expressed in each subcluster displayed in the heatmap in Fig. 1c.**

| No. | Cluster | Gene | No. | Cluster | Gene |
|---|---|---|---|---|---|
| 1 | CHIP1 | *ATP5MC1* | 31 | CHIP3 | *SLC2A3* |
| 2 | CHIP1 | *RPS2* | 32 | CHIP3 | *ANKRD37* |
| 3 | CHIP1 | *SLC25A6* | 33 | CHIP3 | *SLC6A8* |
| 4 | CHIP1 | *RBM3* | 34 | CHIP3 | *FXYD3* |
| 5 | CHIP1 | *HIST1H4C* | 35 | CHIP3 | *ERO1A* |
| 6 | CHIP1 | *TM4SF1* | 36 | CHIP3 | *CKB* |
| 7 | CHIP1 | *SCD* | 37 | CHIP3 | *ZG16* |
| 8 | CHIP1 | *RASGRP2* | 38 | CHIP3 | *NDRG1* |
| 9 | CHIP1 | *ALDH1A1* | 39 | CHIP3 | *SLC11A2* |
| 10 | CHIP1 | *CDK1* | 40 | CHIP3 | *LDHA* |
| 11 | CHIP1 | *MUC13* | 41 | CHIP3 | *HILPDA* |
| 12 | CHIP1 | *RBP2* | 42 | CHIP3 | *APOA4* |
| 13 | CHIP1 | *AREG* | 43 | CHIP3 | *INSIG1* |
| 14 | CHIP1 | *FABP1* | 44 | CHIP3 | *MALAT1* |
| 15 | CHIP1 | *UBE2C* | 45 | CHIP3 | *APOC3* |
| 16 | CHIP2 | *PTTG1* | 46 | TW1 | *ALB* |
| 17 | CHIP2 | *HSP90AA1* | 47 | TW1 | *FGB* |
| 18 | CHIP2 | *HSP90AB1* | 48 | TW1 | *APOH* |
| 19 | CHIP2 | *HMGB1* | 49 | TW1 | *GSTA2* |
| 20 | CHIP2 | *NME1* | 50 | TW1 | *FGG* |
| 21 | CHIP2 | *CDC20* | 51 | TW1 | *RBP4* |
| 22 | CHIP2 | *NCL* | 52 | TW1 | *PHGR1* |
| 23 | CHIP2 | *RANBP1* | 53 | TW1 | *FGA* |
| 24 | CHIP2 | *CCNB1* | 54 | TW1 | *GPC3* |
| 25 | CHIP2 | *PPP1R14B* | 55 | TW1 | *HEPACAM2* |
| 26 | CHIP2 | *GAL* | 56 | TW1 | *GSTA1* |
| 27 | CHIP2 | *DDX21* | 57 | TW1 | *APOE* |
| 28 | CHIP2 | *TUBA1B* | 58 | TW1 | *TPD52L1* |
| 29 | CHIP2 | *HMGB2* | 59 | TW1 | *SELENOP* |
| 30 | CHIP2 | *MYC* | 60 | TW1 | *DKK1* |

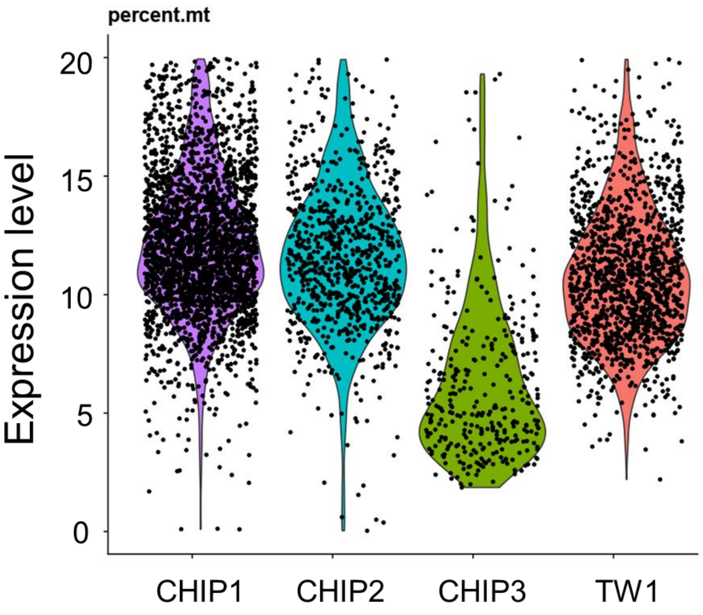


**Extended Data Fig. 1. Mitochondrial gene expression (MT %) of each cell cluster.** Cells in CHIP3 showed significantly lower MT (%) expression than other subclusters.

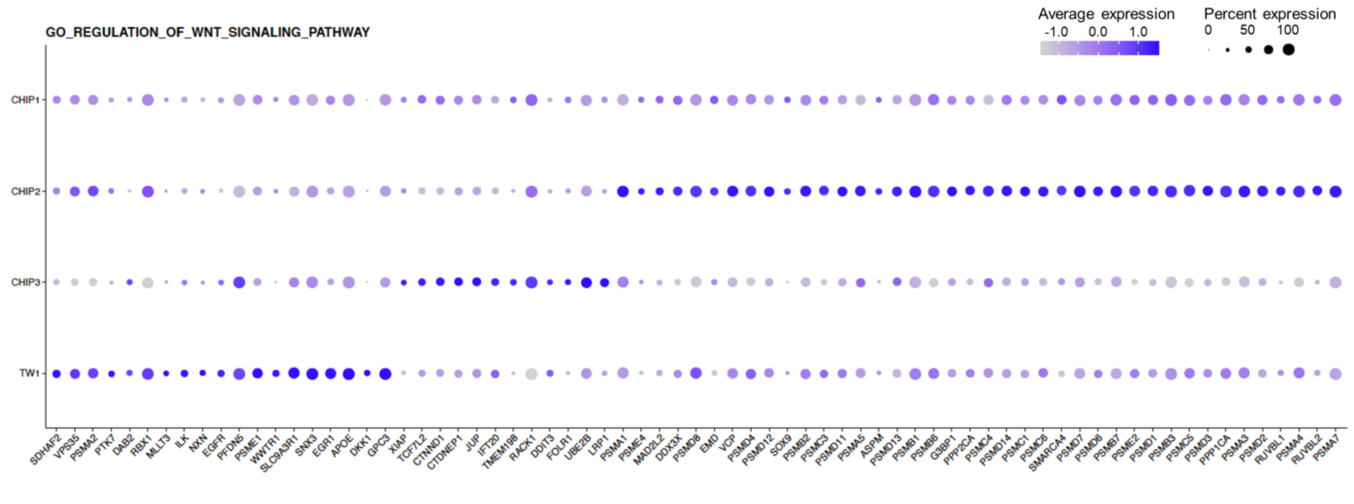


**Extended Data Fig. 2. Single-cell profile of Caco-2 cells associated with the Gene Ontology of "Regulation of Wnt signaling pathway".** Expression of genes involved in Wnt signaling pathway in each subcluster showed heterogenous pattern of cell clusters. Cells in CHIP2 shows the largest number of genes that have upregulated expression.

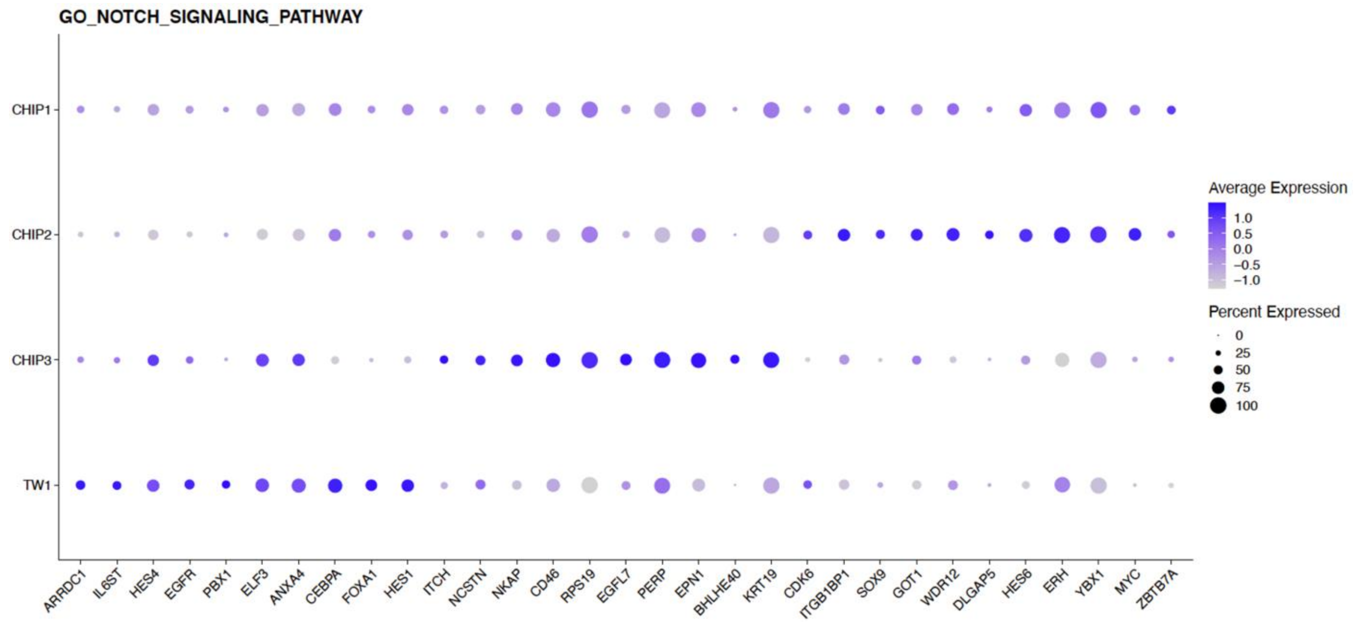


**Extended Data Fig. 3. Single-cell profile of Caco-2 cells associated with the Gene Ontology of "Notch signaling pathway".** CHIP clusters demonstrated the heterogenous expression of genes involved in Notch signaling pathway. Similar to the investigation of gene expressions involved in Wnt signaling pathway, cells in CHIP1 showed decreased expressions.

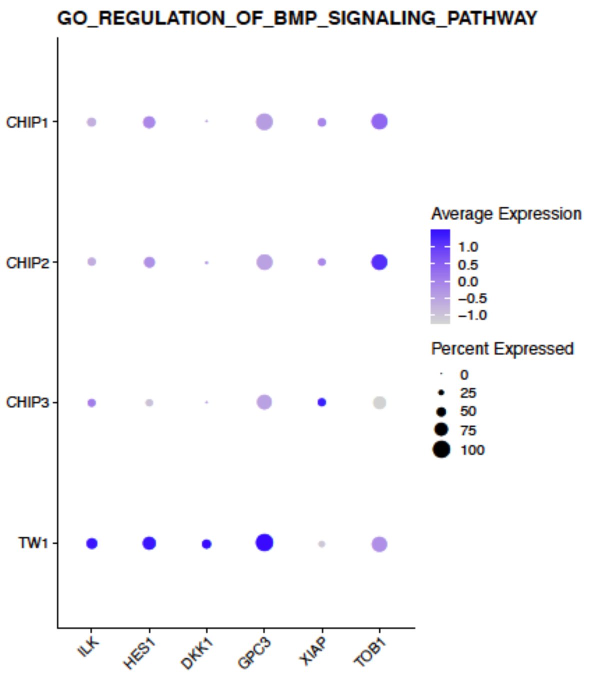


**Extended Data Fig. 4. Expression of genes involved in the regulation of bone morphogenic protein (BMP) pathway.** Cells in TW1 showed increased expression of genes pertinent to the regulation of BMP signaling pathway. BMP signaling is known to negatively regulate the epithelial growth, implying the cells in TW1 are inhibited with cell growth.

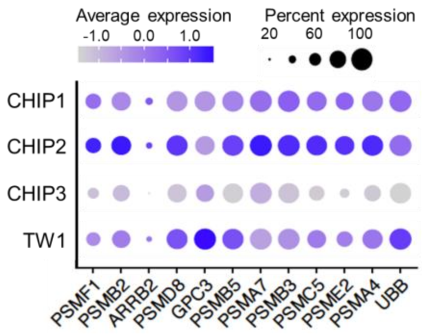


**Extended Data Fig. 5. Heterogenous expression of genes involved in planar cell polarity.** Among the CHIP clusters, cells in CHIP2 demonstrated the strongest expression of genes related to planar cell polarity, whereas cell in CHIP3 showed significantly downregulated expression.

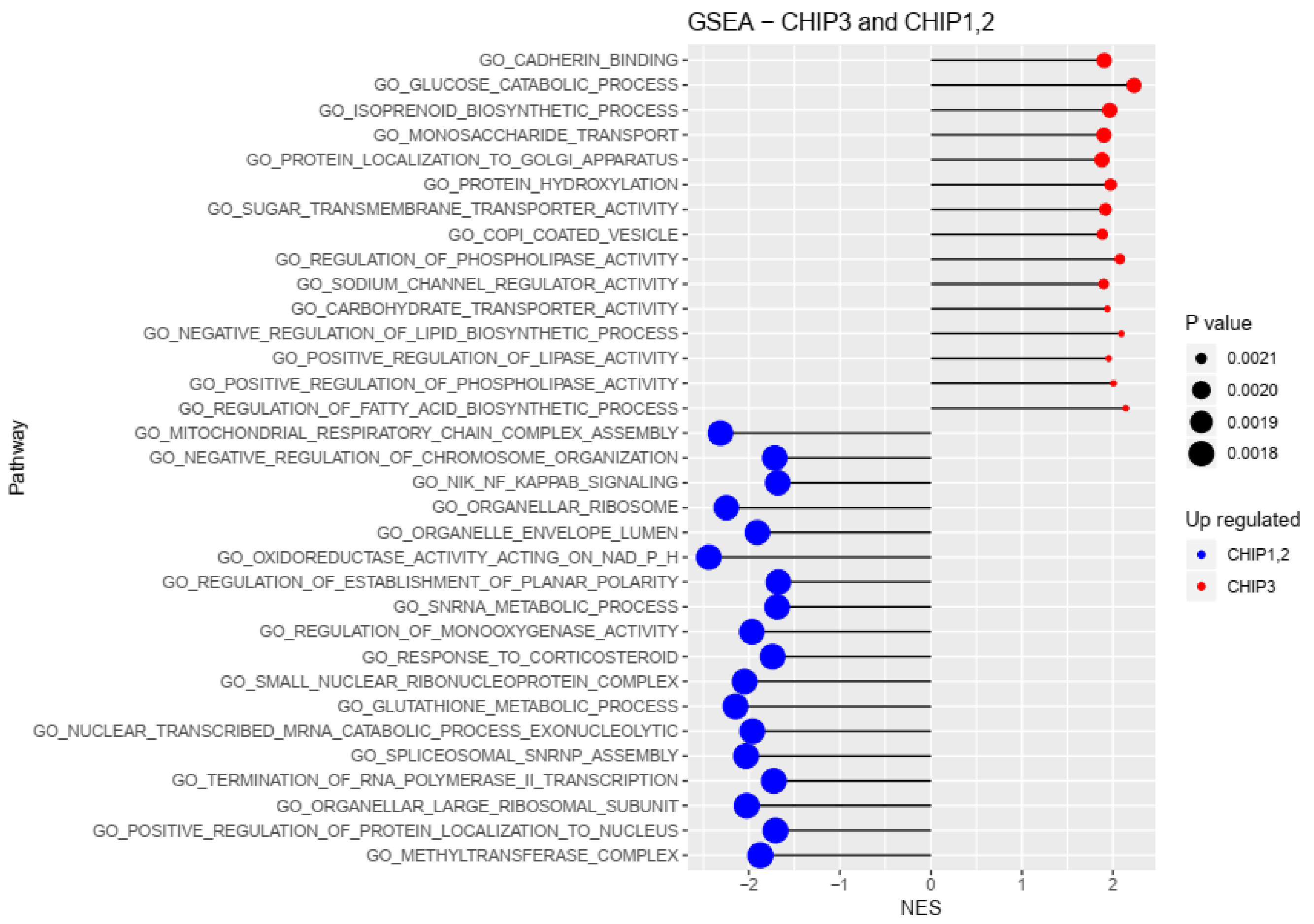


**Extended Data Fig. 6. Most significantly variable pathways in GSEA comparing CHIP3 versus CHIP1 and CHIP2.** Pathways that are significantly upregulated in CHIP3 were highlighted in red, whereas pathways that are upregulated in CHIP1 and 2 were highlighted in blue.

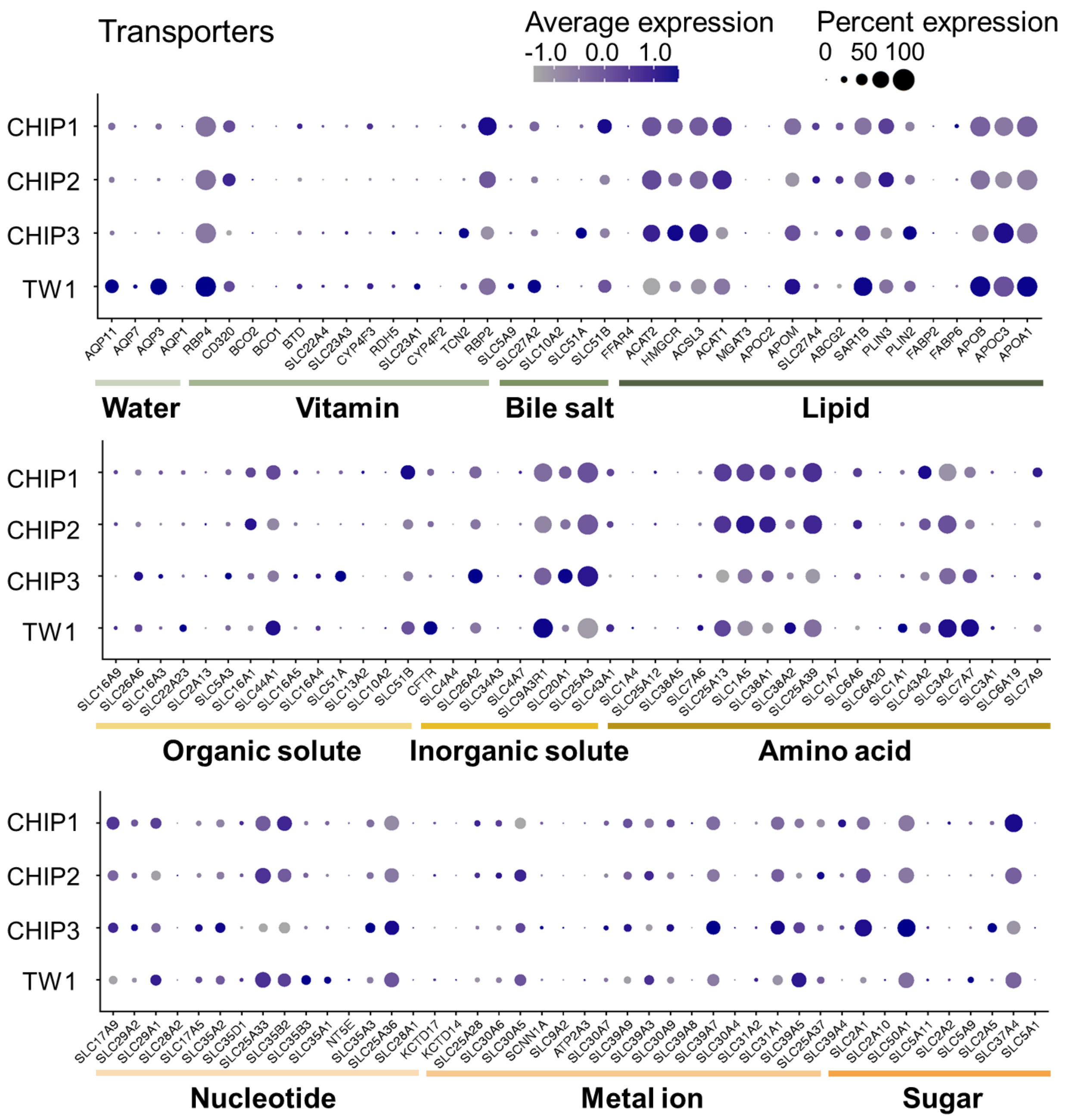


**Extended Data Fig. 7. Expression of genes regulating molecular transport.** Genes encoding transporters of water, vitamin, bile salt, lipid, organic solute, inorganic solute, amino acid, nucleotide, metal ion, and sugar showed heterogenous reprogrammed expressions between clusters.

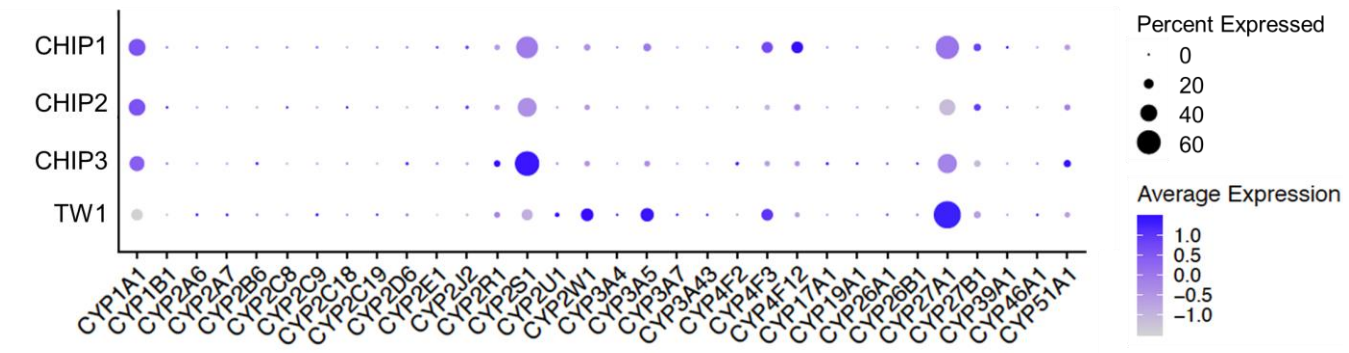


**Extended Data Fig. 8. Expression of genes in cytochrome P450 family.** Genes encoding cytochrome P450 enzymes showed reprogrammed expressions in CHIP clusters.
